# Alveolar macrophage lipid burden correlates with clinical improvement in patients with Pulmonary Alveolar Proteinosis

**DOI:** 10.1101/2022.08.01.502377

**Authors:** Elinor Lee, Kevin J. Williams, Cormac McCarthy, James P. Bridges, Elizabeth F. Redente, Thomas Q. de Aguiar Vallim, Tisha Wang, Elizabeth J. Tarling

## Abstract

Pulmonary alveolar proteinosis (PAP) is a life-threatening rare lung syndrome characterized by the accumulation of surfactant and lipid-loaded macrophages within the alveoli for which there is no cure and no approved therapies. The clinical diagnosis of PAP, often made by invasive lung biopsies and/or cytology of bronchoalveolar lavage fluid does not identify the underlying cause of disease. In addition, no biomarkers exist to inform prognosis or therapeutic options in PAP. We now report on the use of comprehensive mass spectrometry to profile and define the lipid signature of alveolar macrophages obtained from PAP patients. In addition, we quantify how these macrophage-associated lipids change during clinical treatment. Our studies demonstrate that clinical improvement in treated PAP patients is associated with a decrease in total lipid content, indicating that levels of these macrophage-associated lipids correlate with the severity of the disease.

## Introduction

Pulmonary alveolar proteinosis (PAP) is a rare lung disorder with no cure or approved therapies (1). The pathogenesis of this syndrome is heterogenous with various biochemical defects that lead to the accumulation of surfactant within the alveoli. It is categorized into primary, secondary, and congenital etiologies (2). The most common form of disease is primary PAP that occurs either from high levels of neutralizing antibodies to GM-CSF or from hereditary mutations in the GM-CSF receptor (3, 4). Congenital PAP is the rarest disease type and is caused by mutations in genes that are essential for surfactant production (2). Secondary PAP is associated with underlying diseases that secondarily affect alveolar macrophages such as hematologic malignances, immunologic syndromes, infections, or toxic inhalational exposures (1). Approximately 90% of patients have primary autoimmune PAP (aPAP), which is characterized by the presence of autoantibodies against GM-CSF, leading to the disruption of GM-CSF signaling in alveolar macrophages (1). GM-CSF has been identified as a key pulmonary hormone that mediates alveolar macrophage maturation and self-renewal and also regulates population size (5, 6). In the absence of GM-CSF signaling, alveolar macrophages are impaired in their ability to regenerate and mature and also catabolize surfactant (6). Mice deficient in GM-CSF (*Csf2*^*-/-*^) or its receptor (*Csf2ra*^*-/-*^ and *Csf2rb*^*-/-*^) develop PAP-like pulmonary histopathology and are used as pre-clinical mouse models to study PAP (7-9). Current treatment is limited and focused on managing symptoms and treating disease complications. The standard of care is whole lung lavage (WLL), which is invasive, not widely available, and often only transiently effective; however, some experimental therapies including administration of inhaled GM-CSF have shown recent promise (2, 10).

Surfactant is comprised of 85% phospholipids, the majority of which is saturated dipalmitoyl phosphatidylcholine (DPPC; PC16:0/16:0), 10% neutral lipids (mostly cholesterol ester), and 5% proteins (11). Surfactant homeostasis is maintained by the actions of alveolar type II epithelial cells and alveolar macrophages. While alveolar type II cells synthesize and secrete surfactant, about 20-30% of surfactant is catabolized by alveolar macrophages, while the majority is recycled or catabolized by type 2 epithelial cells (12, 13). It was originally proposed that PAP was the result of impaired catabolism of phospholipids in alveolar macrophages (14-16). More recent studies comparing WT, *Csf2rb*^*-/-*^, and *Csf2*^*-/-*^ mice have demonstrated that although *Csf2*^*-/-*^ mice have significantly higher levels of both saturated PC and cholesterol in surfactant, the relative proportion of surfactant cholesterol to total phospholipids is elevated, while the relative proportion of saturated to total phospholipids remained comparable in both groups (17). In these same studies it was also shown that it is the cholesterol content of surfactant that promotes development of disease (17).

We and others have also demonstrated that alveolar macrophages isolated from either PAP patients or *Csf2*^*-/-*^ and *Csf2rb*^*-/-*^ mice have increased levels of cholesterol in addition to changes in the level of triglycerides, free fatty acids, and phospholipid species when compared to macrophages obtained from healthy counterparts (17, 18). It has also been reported that alveolar macrophages isolated from PAP patients or mouse models of PAP have significantly lower expression of the ATP binding cassette transporters A1 and G1 (ABCA1 and ABCG1, respectively) and peroxisome proliferator-activated receptor gamma (PPARγ), important mediators of cholesterol and lipid efflux from macrophages (19-24). Furthermore, we recently reported that treatment of a subset of PAP patients with statin, a drug that targets HMG-CoA reductase leading to reductions in plasma LDL cholesterol levels, markedly reduced pulmonary abnormalities and improved PAP disease (18). Taken together, these data are consistent with the idea that alterations in cholesterol homeostasis have a significant role in the pathogenesis of PAP (17, 18).

Although the lipid composition of bronchoalveolar lavage fluid from patients with PAP was recently described, surprisingly the detailed lipidome of alveolar macrophages obtained from PAP patients remains unknown (25). Given the key role of alveolar macrophages in cholesterol metabolism and the catabolism of surfactant lipids, we hypothesized that defining the lipid composition of these cells might provide new mechanistic insight into PAP and identify novel therapeutic targets. We also hypothesized that lipid composition changes could correlate with a patient’s clinical course, which remains an area of significant unmet need in the PAP community. Here, we report on studies that utilized unbiased comprehensive mass spectrometry to quantify and characterize the lipidome of alveolar macrophages from nonPAP and PAP patients. Further, we report on the lipid profile of alveolar macrophages from two PAP patients during their clinical course. Based on these data, we have identified a relationship between specific macrophage lipid species and the severity of PAP disease.

## Materials and Methods

### Patient selection and ethical approval

All patients included in this study were male and female adults who have underlying nonPAP pulmonary disease or autoimmune pulmonary alveolar proteinosis (PAP) and are being treated at the University of California, Los Angeles. Ages ranged from 26 to 72 years old. The institutional review board of the University of California, Los Angeles approved this study. All human participants provided written informed consent.

### Collection of human BAL fluid and isolation alveolar macrophages

Human BAL fluid was collected from discarded material of nonPAP patients undergoing bronchoscopy with diagnostic bronchoalveolar lavage (BAL) or PAP patients undergoing therapeutic WLL. NonPAP patients had bronchoscopies with BAL performed to evaluate for non-PAP related issues, including infections and airway complications. None of these patients went on to develop secondary PAP. Alveolar macrophages were isolated by centrifugation at 200 g for 5 minutes. Cells were washed and resuspended with Ammonium-Chloride-Potassium (ACK) lysing buffer to lyse red blood cells. Cells were counted and aliquoted at 2×10^6^ or 5×10^6^ cells in 200 μL of phosphate-buffered solution (PBS) before being stored at - 80°C. Macrophage purity was >95% as previously described (21, 24, 26). For frozen samples, cells were frozen immediately at -80°C and counted after being thawed and prior to lipid extraction. The duration of the cells in the freezer ranged from 0.1-8 years before analysis. Long-term stability of the lipids in human lavage fluid for these lengths of time has been previously described (25).

### Lipid extraction and analysis for lipidyzer

A modified Bligh and Dyer extraction was performed on human alveolar macrophages (27). For alveolar macrophages, we used 5×10^6^ cells in 200 μL of PBS. Prior to the biphasic extraction, a 13 lipid class Lipidyzer Internal Standard Mix (AB Sciex, 5040156) was added to each sample. Two successive extractions were done, and the resulting organic layers were pooled and dried down in a Genevac EZ-2 Elite. Lipid samples were then resuspended in 1:1 methanol/dichloromethane with 10 mM ammonium acetate and transferred to robovials (Thermo, 10800107) for analysis.

Samples were analyzed on the Sciex Lipidyzer Platform for targeted quantitative measurement of over 1,100 lipid species across 13 classes including cholesterol esters (CE), ceramides (Cer d18:1), dihydroceramides (Cer d18:0), diacylglycerols (DG), free fatty acids (FFA), hexosylceramides (HexCER), lactosylceramides (LacCER), lysophosphatidylcholines (LPC), lysophosphatidylethanolamines (LPE), phosphatidylcholines (PC), phosphatidylethanolamines (PE), sphingomyelins (SM), and triacylglycerols (TG). The quantification processing has been well described previously (28, 29). Differential Mobility Device on the Lipidyzer was tuned with SelexION tuning kit (Sciex, 5040141). The instrument settings, tuning settings, and multiple reaction monitoring (MRM) list are available upon request. The data was analyzed on the Lipidyzer software with the quantitative values being normalized to cell counts.

### Processing and analysis of samples for gas chromatography/mass spectrometry (GC-MS)

For alveolar macrophages, we used 2×10^6^ cells in 200 μL of PBS to perform acid methanolysis with methanol, toluene, and hydrochloric acid (HCl). Standards were created from a stock cholesterol standard (Avanti Polar Lipids, 700000P-100mg) and diluted with hexane. Prior to adding the acid methanolysis mix, a stigmastanol internal standard lipid was added to each sample and standard. Once the mix was added, the samples were incubated at 45°C for about 8-12 hours. Then, lipids were extracted by adding 1:1 0.4M NaCl and hexane, mixing, spinning, and transferring the top layer to new tubes. This then underwent cholesterol derivatization via sialylation reaction with pyridine (Sigma, 270970-100 mL) and N,O-Bis(trimethylsilyl)trifluoroacetamide with trimethylchlorosilane (BSTFA+TMCS; Supelco, 33155-4) before being run on the GC-MS machine to measure total cholesterol. The instrument settings and tuning settings are available upon request. The data were analyzed on the MassHunter software with quantitative values being normalized to cell counts.

### Statistical analysis

Results are represented as mean ± SEM. Statistical analysis was done with GraphPad Prism 8 software. Paired t-tests were performed for comparison of two time points in an individual’s clinical course. Two-way ANOVA followed by Bonferroni correction was done for comparison of three groups. N and P values are reported in the figure legends. P value <0.05 was used to indicate statistical significance.

## Results

### Alveolar macrophages from PAP patients show broad changes in lipid homeostasis

In order to define the lipid signature in alveolar macrophages, we performed bronchoscopy with bronchoalveolar lavage (BAL) on nonPAP (Table 1) and whole lung lavage (WLL) on PAP patients (Table 2). Lipids were extracted from isolated macrophages and analyzed by mass spectrometry.

**Table 1:** Baseline characteristics of 11 individual nonPAP and 4 individual PAP patients.

| Characteristics | Non PAP 1 | Non PAP 2 | Non PAP 3 | Non PAP 4 | Non PAP 5 | Non PAP 6 | Non PAP 7 | Non PAP 8 | Non PAP 9 | Non PAP 10 | Non PAP 11 |
| --- | --- | --- | --- | --- | --- | --- | --- | --- | --- | --- | --- |
| Age (years old) | 71 | 72 | 60 | 49 | 63 | 65 | 59 | 35 | 68 | 63 | 48 |
| Sex | Male | Male | Male | Male | Male | Male | Female | Female | Male | Female | Male |
| Race | White | Black | White | White | Asian | White | White | Asian | White | Hispanic | Black |
| BMI | 18.5-24.9 | 25-29.9 | 25-29.9 | 25-29.9 | 25-29.9 | 25-29.9 | 18.5-24.9 | 25-29.9 | 25-29.9 | 25-29.9 | 30-39.9 |
| Diagnosis | Lung nodules | Lung cancer | Bronchi-ectasis | Tracheal stenosis | Lung cancer | Met head and neck cancer | Familial ILD | CTD-ILD | COPD | RA-ILD | Sarcoid-osis |
| Lung transplant status | No | No | No | No | No | No | Post | Post | Post | Post | Post |
| Smoking status | Former | Former | Never | Former | Former | Former | Never | Never | Former | Never | Never |
| Co-morbidities |  |  |  |  |  |  |  |  |  |  |  |
| Hyperlipidemia | Yes | Yes | No | No | No | Yes | No | No | Yes | Yes | Yes |
| Hypertension | Yes | No | No | No | No | Yes | Yes | No | Yes | No | Yes |
| Diabetes | No | No | No | No | No | No | No | No | Yes | No | No |

**Table 2:** Baseline characteristics of 8 individual autoimmune PAP patients on no or various experimental therapies.

| Characteristics | PAP 1 | PAP 2 | PAP 3 | PAP 4 | PAP 5 | PAP 6 | PAP 7 | PAP 8 |
| --- | --- | --- | --- | --- | --- | --- | --- | --- |
| Age (years old) | 42 | 54 | 50 | 63 | 29 | 59 | 34 | 26 |
| Sex | Male | Female | Male | Female | Male | Male | Female | Female |
| Race | White | Black | Black | Hispanic | White | Asian | Hispanic | Black |
| BMI | 18.5-24.9 | ≥40 | 25-29.9 | 30-39.9 | 25-29.9 | 18.5-24.9 | 18.5-24.9 | 25-29.9 |
| Imaging- presence of pulmonary fibrosis | No | No | No | No | No | Yes | Yes | Yes |
| Treatment (apart from WLL) | None | Statin | Rituximab, plasmapheresis, statin, pioglitazone | Statin | None | Statin | Inhaled GM-CSF | None |
| Smoking status | Never | Never | Never | Never | Never | Never | Never | Never |
| Co-morbidities |  |  |  |  |  |  |  |  |
| Hyperlipidemia | No | Yes | Yes | Yes | No | Yes | No | No |
| Hypertension | No | Yes | Yes | Yes | No | Yes | No | Yes |
| Diabetes | No | No | Yes | Yes | No | Yes | No | No |

Alveolar macrophages from PAP patients demonstrated an overall increase in total lipid content compared to nonPAP patients (Figure 1A). The predominant lipid class recovered in both nonPAP and PAP alveolar macrophages was phosphatidylcholine (PC) (Figure 1A,B; teal). The individual lipid profiles of nonPAP patients displayed broad heterogeneity in terms of lipid composition (Figure 1C and Supplemental Figure 1A), possibly due to individual variability or the variety of pulmonary diseases in these patients (Table 1). Overall, we see a marked difference in the lipid composition of the alveolar macrophages from PAP patients compared to nonPAP patients (Figure 1B and Supplemental Figure 1A,B). PC and PE species represent 46% and 18%, respectively, of the total lipid in nonPAP alveolar macrophages, whereas in PAP alveolar macrophages PC represents almost 75% of the total lipid and PE represents only 2% (Figure 1B). In almost all of the nonPAP patients, the major lipids recovered from the macrophages were phosphatidylcholine (PC) and phosphatitidyethanolamine (PE) (Figure 1B,C; teal and orange, respectively). These cells also contained lower abundance of other major lipid classes, which included free fatty acids (FFA; green), sphingomyelin (SM; salmon), triglyceride (TG; yellow), and cholesterol esters (CE; red) (Figure 1B,C).

**Figure 1.**
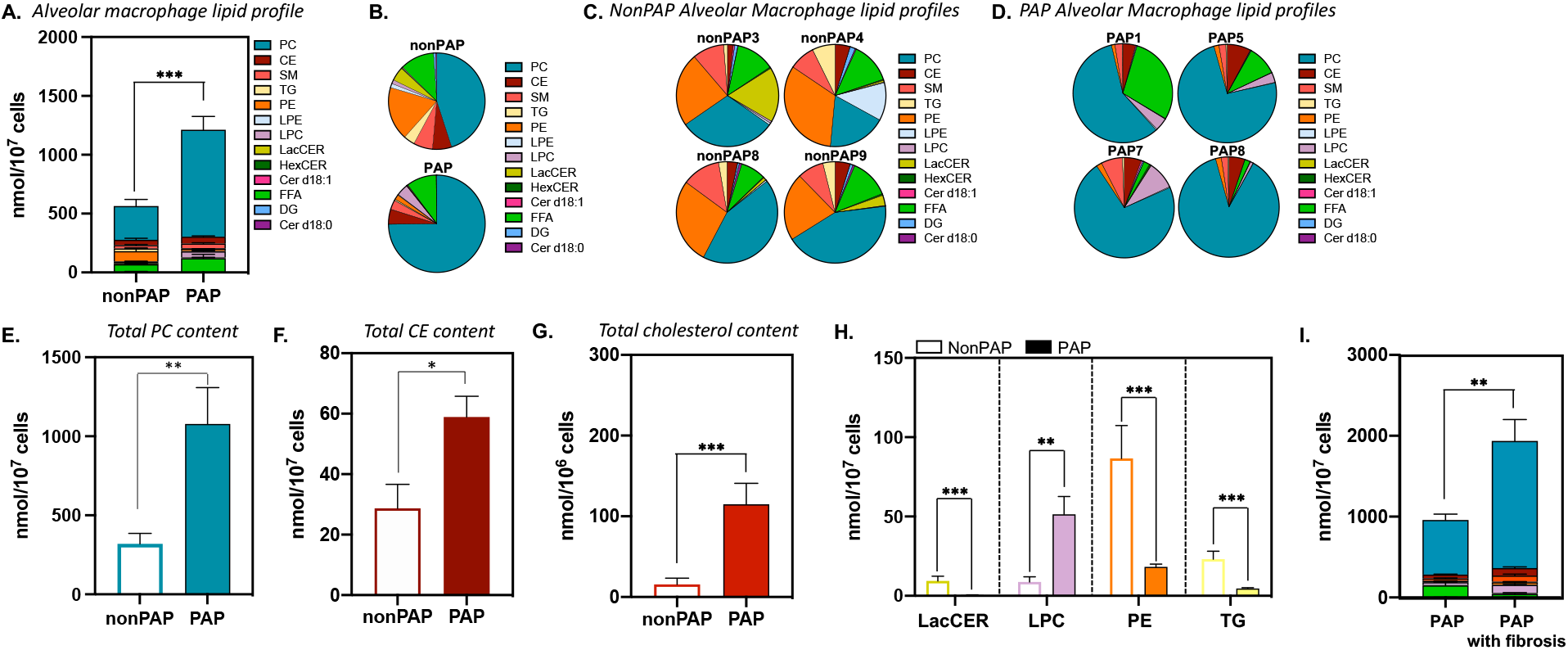
High-resolution unbiased lipidomic profiling reveals broad changes in lipid metabolism in PAP alveolar macrophages. (A) Quantitative measurement of lipid class profile in alveolar macrophages of nonPAP patients vs PAP patients. (B) Compositional lipid analysis of alveolar macrophages from PAP patients compared to nonPAP patients. (C-D) Quantitative compositional analysis of lipid classes in alveolar macrophages of individual nonPAP patients (C) and PAP patients (D). (E-F) Total PC (E) and CE (F) content in nonPAP vs PAP patients. (G) Total cholesterol content measured by GC/MS in alveolar macrophages of PAP patients compared to nonPAP patients. (H) Total LacCer, LPC, PE, and TG content in alveolar macrophages of nonPAP and PAP patients. (I) Quantitative measurement of lipid class profile in alveolar macrophages of PAP patients and PAP patients with fibrosis. All samples are run in duplicate or triplicate. Data are mean ± SEM. CE = cholesterol esters; Cer d18:1= ceramides; Cer d18:0= dihydroceramides; DG= diacylglycerols; FFA= free fatty acids; HexCER= hexosylceramides; LacCER= lactosylceramides; LPC= lysophosphatidylcholines; LPE= lysophosphatidylethanolamines; PC= phosphatidylcholines; PE= phosphatidylethanolamines; SM= sphingomyelins; TG= triacylglycerols. Significance was determined at *p*<0.05 by Student’s *t*-test. * *p*<0.05, ** *p*<0.01, *** *p*<0.001.

In all PAP patient alveolar macrophage samples the predominant lipid species recovered was PC (Figure 1B,D), with levels increased by 3-fold in PAP alveolar macrophages compared to nonPAP alveolar macrophages (Figure 1E). Absolute levels of CE were significantly higher in PAP alveolar macrophages (Figure 1F). Consistent with our previously published data (18), total cholesterol content as measured by gas chromatography-mass spectrometry (GC-MS) was significantly elevated 7-fold in PAP alveolar macrophages compared to nonPAP alveolar macrophages (Figure 1G). Lysophosphatidylcholine (LPC; lilac) levels were significantly elevated 11-fold in PAP alveolar macrophages compared to nonPAP alveolar macrophages, while lactosyl ceramides (LacCER), PE, and TG were all significantly decreased by 13.8-fold, 5.3-fold, and 4.8-fold, respectively (Figure 1H). We observed smaller increases in d18:1 sphingosine ceramides (Cer d18:1) and free fatty acids (FFA), with no significant differences in diacylglycerol (DG), d18:0 ceramides, hexosyl ceramides (HexCer), LPE, and sphingomyelin (SM) (Supplemental Figure 1C). When we specifically compare PAP patients with fibrosis (PAP6, PAP7, and PAP8) to PAP patients without fibrosis, we find that PAP patients with fibrosis have significantly increased lipid content within their alveolar macrophages (Figure 1I). There are specific increases in CE (Supplemental Figure 1D) that are accompanied by increases in Cer d18:1, HexCer, LPC, LPE, and TG (Supplemental Figure 1E). Taken together, these data suggest that, in addition to the previously observed increases in PC and cholesterol, these broad changes in lipid metabolism in alveolar macrophages from PAP patients may have important implications for disease pathogenesis and progression.

Although the major surfactant lipid in both nonPAP and PAP macrophages is PC, we identified significant differences in PC species composition in alveolar macrophages between PAP and nonPAP patients (Figure 2A,B) (25, 30). In nonPAP alveolar macrophages, PC containing 16:0/16:0 or 16:0/18:1 fatty acids represent approximately 27% and 20% of the total cellular PC, respectively (Figure 2B). In contrast, in PAP alveolar macrophages, PC16:0/16:0 represents almost 50% and PC16:0/18:1 represents only 13% of the total cellular PC (Figure 2B). Furthermore, both the saturated (Figure 2C) and unsaturated (Figure 2D) PC content were significantly increased in PAP macrophages, compared to nonPAP macrophages. Similar to what we observed with PC species composition (Figure 2B), the composition of CE species was significantly altered (Figure 2F). The most abundant CE species in nonPAP alveolar macrophages was CE18:2, whereas in PAP alveolar macrophages the predominant CE species were CE16:0 and CE18:0 (Figure 2F). Saturated CE16:0 and unsaturated CE18:2 represent approximately 15% and 33% of the total CE, respectively, in nonPAP alveolar macrophages (Figure 2F). In contrast, in PAP alveolar macrophages CE16:0 and CE18:2 represent 30% and 20% of total CE species, respectively (Figure 2F). These changes are reflected by a significant increase in total saturated CE content in PAP alveolar macrophages but no significant difference in unsaturated content (Figure 2G,H). Following our observations that additional lipids are also altered in PAP alveolar macrophages (Figure 1I), we performed further compositional analysis (Supplemental Figure 2). There were striking changes in diacylglycerol (DG), lactosylceramide (LacCer), LPC, PE and SM composition (Supplemental Figure 2). Together, these data demonstrate significant changes in alveolar macrophage lipid composition in addition to the well documented accumulation of surfactant PC in patients with PAP.

**Figure 2.**
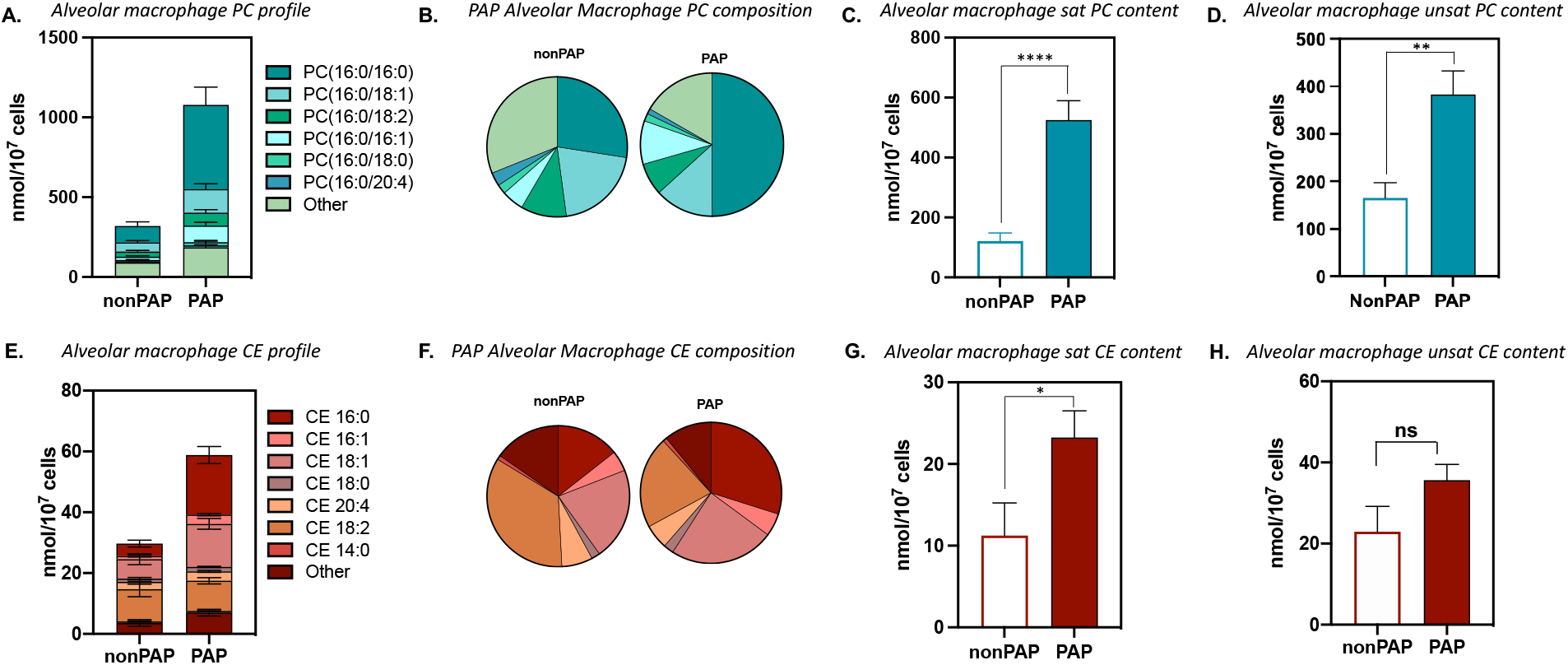
PAP alveolar macrophages display a significant shift in lipid compositional content. (A-B) Absolute quantification (A) and compositional analysis (B) of the most abundant PC species, including PC(16:0/16:0), PC(16:0/16:1), PC(16:0/18:0), PC(16:0/18:1), PC(16:0/18:2), and PC(16:0/20:4), in alveolar macrophages from PAP patients compared to nonPAP patients. (C-D) Quantitative measurement of saturated (C) and unsaturated (D) PC content in alveolar macrophages from nonPAP vs PAP patients. (E-F) Absolute quantification (E) and compositional analysis (F) of the most abundant CE species, including CE(14:0), CE(16:0), CE(16:1), CE(18:0), CE(18:1), CE(18:2), and CE(20:4), in alveolar macrophages from PAP patients compared to nonPAP patients. (G-H) Quantitative measurement of saturated (G) and unsaturated (H) CE content in alveolar macrophages from nonPAP vs PAP patients. All samples are run in duplicate or triplicate. Data are mean ± SEM. CE = cholesterol esters; PC= phosphatidylcholines. Significance was determined at *p*<0.05 by Student’s *t*-test. * *p*<0.05, ** *p*<0.01, **** *p*<0.0001..

### Lipid profile of alveolar macrophages changes during patients’ clinical courses

PAP patients often display variable disease progression ranging from progressive deterioration to stable, yet unremitting disease. However, the reasons underlying these marked differences in clinical course are not well understood. Further, there are no reliable biomarkers of a patient’s disease progression or response to therapy. Because PAP is a disease of lipid accumulation, we hypothesized that the lipid profile (composition and/or total lipid level) of alveolar macrophages would correlate with a patient’s disease progression or regression. Here, we provide examples of two PAP patients, PAP 1 and PAP 7 (Table 2). The first patient, PAP 1, is an otherwise healthy 42 year old male, who was diagnosed with aPAP in early 2014 (Figure 3). He required 2-3 WLL per year due to dyspnea before being initiated on inhaled GM-CSF in February 2016. After receiving inhaled GM-CSF for approximately 9 months, his clinical status improved markedly, and he has not required further WLL to date (Figure 3A). His clinical improvement is evidenced by the decreased ground glass opacities (red asterisk) and septal thickening (red arrowhead) on imaging between his diagnostic CT in 2014 prior to treatment (Figure 3B, No Tx) and his 2018 CT after receiving multiple WLL and inhaled GM-CSF for 33 months (Figure 3B, WLL+Tx).

**Figure 3.**
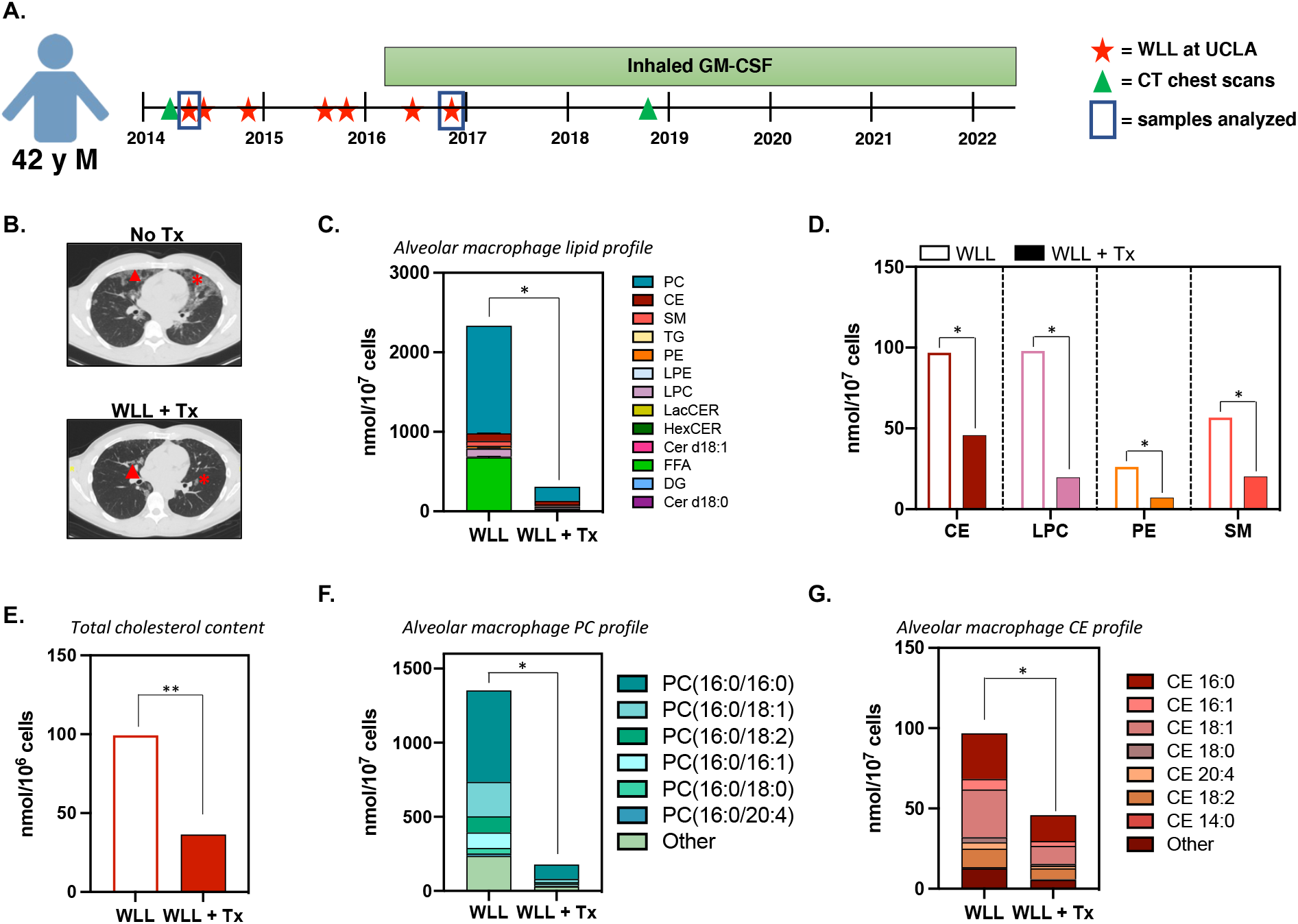
Decreased alveolar macrophage lipid burden is associated with clinical improvements in PAP disease in response to therapy. (A) Clinical course of a 42 year old patient with disease improvement after being initiated on inhaled GM-CSF. Red stars indicate whole lung lavages that were performed at UCLA. Green triangles denote when CT chest scans were taken. Blue boxes represent whole lung lavages used for analysis in (C-G). (B) CT chest scans of the patient at the time of presentation to UCLA (March 2014; No Tx) and after whole lung lavage and inhaled GM-CSF therapy (November 2018; WLL+Tx). Red stars indicate ground glass opacities, and red arrowheads indicate septal thickening. (C) Quantitative measurement and compositional analysis of total lipid content in alveolar macrophages of the patient before (WLL) and after initiation of inhaled GM-CSF (WLL+Tx). (D) Total CE, LPC, PE, and SM content in alveolar macrophages of the patient before (WLL) and after initiation of inhaled GM-CSF (WLL+Tx). (E) Total cholesterol content measured by GC/MS before (WLL) and after patient has been on inhaled GM-CSF (WLL+Tx). (F-G) Absolute quantification of the most abundant PC (F) and CE (G) species in alveolar macrophages of the patient before (WLL) and after being placed on inhaled GM-CSF (WLL+Tx). Samples are run in triplicate. Data are mean ± SEM. Statistical significance determined by student’s t test. * p<0.05, ** p<0.01, *** p<0.001, **** p<0.0001. WLL= whole lung lavage; Tx = treatment, which includes inhaled GM-CSF. CE = cholesterol esters; Cer d18:1= ceramides; Cer d18:0= dihydroceramides; DG= diacylglycerols; FFA= free fatty acids; HexCER= hexosylceramides; LacCER= lactosylceramides; LPC= lysophosphatidylcholines; LPE= lysophosphatidylethanolamines; PC= phosphatidylcholines; PE= phosphatidylethanolamines; SM= sphingomyelins; TG= triacylglycerols.

Alveolar macrophages were obtained in 2014 shortly after diagnosis and after initiation on inhaled GM-CSF (Figure 3A; blue boxes). Unbiased lipidomic profiling of the patient’s alveolar macrophages in 2014, shortly after his diagnosis, revealed that the most abundant lipid classes were PC, free fatty acids, and CE. As additional evidence of the effectiveness of inhaled GM-CSF therapy, the total lipid content and overall levels of all major lipid classes decreased after 10 months (Figure 3C-D). These data demonstrate that the macrophages present in the alveolar space after 10 months of GM-CSF have lipid profiles that are more similar to those seen in nonPAP cells, consistent with normal, or near-normal, macrophage lipid metabolism. Total cholesterol levels as measured by GC-MS were also significantly decreased in his alveolar macrophages 10 months after being on inhaled GM-CSF, indicating improvement in surfactant lipid catabolism (Figure 3E). The overall amount of PC and CE species significantly decreased after the patient was placed on inhaled GM-CSF (Figure 3F-G). Surprisingly, PC and CE species compositions were not drastically altered in alveolar macrophages before and after GM-CSF inhalation (Figure 3F-G), suggesting that changes in overall macrophage lipid burden are more reflective of clinical improvement.

In contrast, the second patient, PAP 7, had a markedly different clinical course (Figure 4A). She is an otherwise healthy 34 year old female, who was diagnosed with aPAP in late 2017. She had two whole lung lavages before developing severe hypoxic respiratory failure requiring admission to the intensive care unit. She was initiated on inhaled GM-CSF but continued to require frequent whole lung lavages (Figure 4A). Because of the severe nature and deterioration of her disease, she received rituximab and was started on a statin (Figure 4A). Despite these interventions, the patient remained short of breath and hypoxic. She was then started on pioglitazone and plasmapheresis with additional doses of rituximab (Figure 4A). The patient eventually underwent a bilateral lung transplant in 2021 due to her declining respiratory status and worsening pulmonary fibrosis in spite of multiple therapies (Figure 4A). A chest CT scan in 2020 after receiving the above therapies and prior to lung transplant (Figure 4B; WLL+Tx) confirmed a lack of clinical improvement with evidence of ongoing diffuse ground glass opacifications (red asterisk) and septal thickening (red arrowhead) with new subpleural fibrotic changes when compared to her CT scan in 2018 after two whole lung lavages (Figure 4B; WLL).

**Figure 4.**
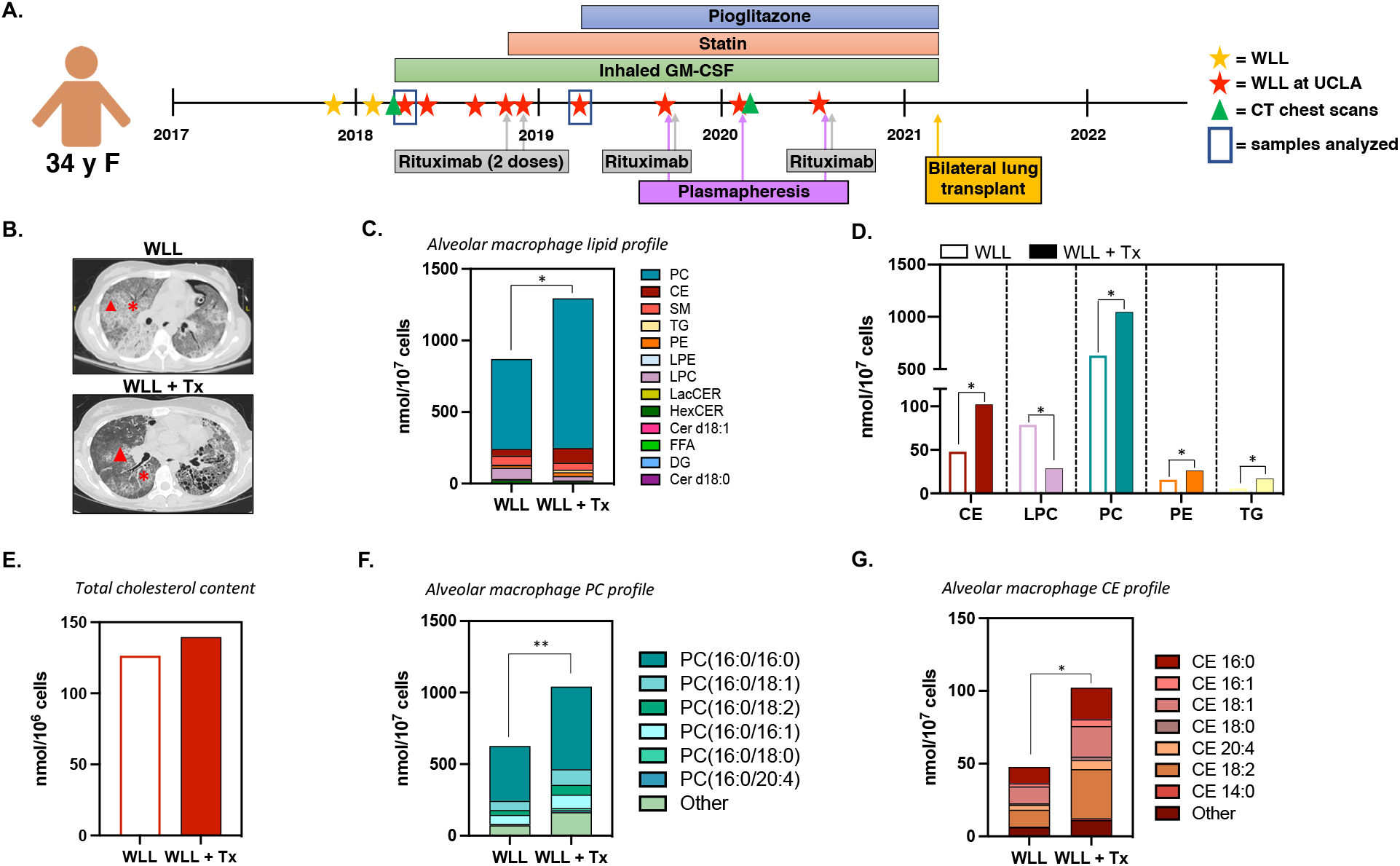
Increased alveolar macrophage lipid burden is associated with worsening, unremitting disease. (a) Clinical course of a 34 year old patient with worsening PAP disease and unresolving hypoxia despite multiple therapies. Stars indicate whole lung lavages. Red stars indicate whole lung lavages that were performed at UCLA. Green triangles denote when CT chest scans were taken and shown in part b. Blue boxes represent when the whole lung lavages from patients were used for analysis in parts d-h. (b) CT chest scans of the patient at the time of presentation to UCLA (April 2018; WLL) and after multiple therapies (January 2020; WLL+Tx). Red stars indicate ground glass opacities, and red arrowheads indicate septal thickening. (C) Quantitative measurement and compositional analysis of total lipid content in alveolar macrophages of the patient before (WLL) and after initiation of inhaled GM-CSF (WLL+Tx). (D) Total CE, LPC, PE, and SM content in alveolar macrophages of the patient before (WLL) and after initiation of inhaled GM-CSF (WLL+Tx). (E) Total cholesterol content measured by GC/MS before (WLL) and after patient has been on inhaled GM-CSF (WLL+Tx). (F-G) Absolute quantification of the most abundant PC (F) and CE (G) species in alveolar macrophages of the patient before (WLL) and after being placed on inhaled GM-CSF (WLL+Tx). Samples are run in triplicate. Data are mean ± SEM. Statistical significance determined by student’s t test. * p<0.05, ** p<0.01, *** p<0.001, **** p<0.0001. WLL= whole lung lavage; Tx = treatment, which includes inhaled GM-CSF. CE = cholesterol esters; Cer d18:1= ceramides; Cer d18:0= dihydroceramides; DG= diacylglycerols; FFA= free fatty acids; HexCER= hexosylceramides; LacCER= lactosylceramides; LPC= lysophosphatidylcholines; LPE= lysophosphatidylethanolamines; PC= phosphatidylcholines; PE= phosphatidylethanolamines; SM= sphingomyelins; TG= triacylglycerols.

Alveolar macrophages were obtained when the patient was first on inhaled GMCSF and after initiation on a statin (Figure 4A; blue boxes). Analysis of the lipid profile of the patient’s alveolar macrophages indicated that the most abundant lipid classes were PC, CE, SM and LPC, which differed from both nonPAP patients and patient PAP 1 (Figure 4C-D compared with Figures 1B and 3C, respectively). Both total PC and CE levels in alveolar macrophages significantly increased as the patient’s disease worsened (Figure 4C-D). In contrast, total macrophage cholesterol levels were only modestly increased (Figure 4E). The fatty acid composition of PC and CE species in the alveolar macrophages was not significantly altered as the patient’s disease progressed; however the total amount of both PC and CE was almost 2-fold increased in just over a year (Figure 4F-G). Consistent with our observations from Patient 1 (Figure 3), these data suggest that changes in overall alveolar macrophage lipid burden, not lipid composition, are the most reflective of a patient’s clinical progress.

## Discussion

PAP is a rare, life-threatening syndrome of lipid accumulation in the alveolar space and particularly within alveolar macrophages (4). Here, we provide the first in-depth study defining the lipidome of human alveolar macrophages in PAP. We show that in comparison to alveolar macrophages recovered from nonPAP patients, PAP-afflicted alveolar macrophages exhibited increased levels of numerous lipids, in addition to the previously reported increases in PC (7-9, 15, 17-19, 22, 25, 31, 32), demonstrating that there are broader changes in lipid metabolism within PAP alveolar macrophages than previously observed using more qualitative analyses. Additionally, we also show that the lipid composition within alveolar macrophages varies considerably for each patient, demonstrating the significant heterogeneity of this disease.

Alveolar macrophages are the main cellular target of the downstream effects of autoantibodies to GM-CSF and loss of GM-CSF signaling, yet lipid profiles of PAP patient alveolar macrophages have not been previously defined, leaving a significant gap in knowledge. Until recently, most lipid profiling studies have focused on lipids within the bronchoalveolar lavage fluid (BALF) in the alveolar space (25). Griese and colleagues reported an overall increase of total lipid concentration within the lavage fluid recovered from the alveolar airspace with free cholesterol being increased the most (60-fold), followed by CE and PC. In their reported sample preparation, whole lung lavage fluid from healthy individuals was centrifuged to remove cellular content, and the remaining lavage fluid was analyzed by direct flow injection electrospray ionization tandem mass spectrometry (ESI-MS/MS). However, in the case of PAP patients, unprocessed lavage fluid including cellular components was utilized for analysis. Because lavage fluid is a complex mixture of surfactant, alveolar macrophages, proteinaceous material, and in some cases other cell types, the inclusion and/or exclusion of these components can significantly affect lipidomic data and interpretation of results. Therefore, due to the differences in sample preparation and methodology, it is not possible to directly compare our findings with those reported by Griese *et al*.

There are currently no quantitative measures of disease burden for PAP patients. In addition, there are no studies on macrophage lipid profiles and disease progression or regression. We noted that the absolute abundance of lipids correlated with disease severity in patients with improving or persistent, worsening clinical disease. Additional studies will be required to determine if the alveolar macrophage lipidome or a specific combination of lipids could be used as a biomarker to determine severity of disease or predict response to therapy.

Alveolar macrophages are the main cell type affected by loss of normal GM-CSF signaling in PAP (4). Our previous study demonstrated that treatment of a subset of PAP patients with statins led to a remarkable improvement in their clinical symptoms, likely by reducing alveolar macrophage cholesterol content (18). Whether statins affect the macrophage CE species in the responsive patients is currently unknown. However, not all patients respond to statin therapy, suggesting that there is a need for additional studies on the lipidome of alveolar macrophages and molecular targets. Our findings support the exciting possibility that other lipid pathways that we identified to be altered in PAP macrophages may also be viable targets for treatment of PAP (17, 18). Elucidation of the lipidome in an increased number of PAP patients, particularly from patients on different therapies, will provide further insight into the pathogenesis of PAP, assist in discovering new biomarkers to predict severity of disease and response to treatment, and identify novel therapeutic targets.

## Acknowledgement/grant support

We would like to thank members of the Tarling and Vallim labs for critical and thoughtful discussion. We thank Dr. Peter Edwards for review and comment of the manuscript. EL was supported by T32HL072752 and U54HL127672. TQdeAV is supported by DK118064. EFR is supported by HL147860 and HL149741. EJT is supported by HL136543, HL152156, DK128952 and UL1TR001881.

## References

1. Trapnell BC, Whitsett JA, Nakata K. Pulmonary alveolar proteinosis. N Engl J Med. 2003;349(26):2527–39.

2. Trapnell BC, Nakata K, Bonella F, Campo I, Griese M, Hamilton J, et al. Pulmonary alveolar proteinosis. Nat Rev Dis Primers. 2019;5(1):16.

3. McCarthy C, Avetisyan R, Carey BC, Chalk C, Trapnell BC. Prevalence and healthcare burden of pulmonary alveolar proteinosis. Orphanet J Rare Dis. 2018;13(1):129.

4. McCarthy C, Carey BC, Trapnell BC. Autoimmune Pulmonary Alveolar Proteinosis. Am J Respir Crit Care Med. 2022;205(9):1016–35.

5. Hamilton JA. GM-CSF in inflammation and autoimmunity. Trends Immunol. 2002;23(8):403–8.

6. Guilliams M, De Kleer I, Henri S, Post S, Vanhoutte L, De Prijck S, et al. Alveolar macrophages develop from fetal monocytes that differentiate into long-lived cells in the first week of life via GM-CSF. J Exp Med. 2013;210(10):1977–92.

7. Shima K, Arumugam P, Sallese A, Horia Y, Ma Y, Trapnell C, et al. A Murine Model of Hereditary Pulmonary Alveolar Proteinosis Caused by Homozygous Csf2ra Gene Disruption. Am J Physiol Lung Cell Mol Physiol. 2022.

8. Robb L, Drinkwater CC, Metcalf D, Li R, Kontgen F, Nicola NA, et al. Hematopoietic and lung abnormalities in mice with a null mutation of the common beta subunit of the receptors for granulocyte-macrophage colony-stimulating factor and interleukins 3 and 5. Proc Natl Acad Sci U S A. 1995;92(21):9565–9.

9. Dranoff G, Crawford AD, Sadelain M, Ream B, Rashid A, Bronson RT, et al. Involvement of granulocyte-macrophage colony-stimulating factor in pulmonary homeostasis. Science. 1994;264(5159):713–6.

10. Ohkouchi S, Akasaka K, Ichiwata T, Hisata S, Iijima H, Takada T, et al. Sequential Granulocyte-Macrophage Colony-Stimulating Factor Inhalation after Whole-Lung Lavage for Pulmonary Alveolar Proteinosis. A Report of Five Intractable Cases. Ann Am Thorac Soc. 2017;14(8):1298–304.

11. Lopez-Rodriguez E, Gay-Jordi G, Mucci A, Lachmann N, Serrano-Mollar A. Lung surfactant metabolism: early in life, early in disease and target in cell therapy. Cell Tissue Res. 2017;367(3):721–35.

12. Goss V, Hunt AN, Postle AD. Regulation of lung surfactant phospholipid synthesis and metabolism. Biochim Biophys Acta. 2013;1831(2):448–58.

13. Rider ED, Ikegami M, Jobe AH. Localization of alveolar surfactant clearance in rabbit lung cells. Am J Physiol. 1992;263(2 Pt 1): L201–9.

14. Griese M. Pulmonary surfactant in health and human lung diseases: state of the art. Eur Respir J. 1999;13(6):1455–76.

15. Ikegami M, Ueda T, Hull W, Whitsett JA, Mulligan RC, Dranoff G, et al. Surfactant metabolism in transgenic mice after granulocyte macrophage-colony stimulating factor ablation. Am J Physiol. 1996;270(4 Pt 1): L650–8.

16. Pison U, Wright JR, Hawgood S. Specific binding of surfactant apoprotein SP-A to rat alveolar macrophages. Am J Physiol. 1992;262: L412–7.

17. Sallese A, Suzuki T, McCarthy C, Bridges J, Filuta A, Arumugam P, et al. Targeting cholesterol homeostasis in lung diseases. Sci Report. 2017;7(1):10211–4.

18. McCarthy C, Lee E, Bridges JP, Sallese A, Suzuki T, Woods JC, et al. Statin as a novel pharmacotherapy of pulmonary alveolar proteinosis. Nat Commun. 2018;9(1):3127.

19. Thomassen MJ, Barna BP, Malur AG, Bonfield TL, Farver CF, Malur A, et al. ABCG1 is deficient in alveolar macrophages of GM-CSF knockout mice and patients with pulmonary alveolar proteinosis. J Lipid Res. 2007;48(12):2762–8.

20. Cavelier C, Lorenzi I, Rohrer L, von Eckardstein A. Lipid efflux by the ATP-binding cassette transporters ABCA1 and ABCG1. Biochim Biophys Acta. 2006;1761(7):655–66.

21. Baldan A, Tarr P, Vales CS, Frank J, Shimotake TK, Hawgood S, et al. Deletion of transmembrane transporter ABCG1 results in progressive pulmonary lipidosis. J Biol Chem. 2006;281:29401–10.

22. Bonfield TL, Farver CF, Barna BP, Malur A, Abraham S, Raychaudhuri B, et al. Peroxisome proliferator-activated receptor-gamma is deficient in alveolar macrophages from patients with alveolar proteinosis. Am J Respir Cell Mol Biol. 2003;29(6):677–82.

23. Kennedy MA, Barrera GC, Nakamura K, Baldan A, Tarr P, Fishbein MC, et al. ABCG1 has a critical role in mediating cholesterol efflux to HDL and preventing cellular lipid accumulation. Cell Metab. 2005;1(2):121–31.

24. de Aguiar Vallim TQ, Lee E, Merriott DJ, Goulbourne CN, Cheng J, Cheng A, et al. ABCG1 regulates pulmonary surfactant metabolism in mice and men. J Lipid Res. 2017;58(5):941–54.

25. Griese M, Bonella F, Costabel U, de Blic J, Tran NB, Liebisch G. Quantitative Lipidomics in Pulmonary Alveolar Proteinosis. Am J Respir Crit Care Med. 2019;200(7):881–7.

26. Baldán A, Gomes AV, Ping P, Edwards PA. Loss of ABCG1 results in chronic pulmonary inflammation. J Immunol. 2008;180(5):3560–8.

27. Bligh EG, Dyer WJ. A rapid method of total lipid extraction and purification. Can J Biochem Physiol. 1959;37(8):911–7.

28. Contrepois K, Mahnoudi S, Ubhi BK, Papsdorf K, Hornburg D, Brunet A, et al. Cross-platform comparison of untargeted and targeted lipidomics approaches on aging mouse plasma. Sci Rep. 2018;8(1):17747.

29. Hsieh W, Williams KJ, Su B, Bensinger SJ. Profiling of mouse macrophage lipidome using direct infusion shotgun mass spectrometry. STAR Protoc. 2020;2(1):100235.

30. Veldhuizen R, Nag K, Orgeig S, Possmayer F. The role of lipids in pulmonary surfactant. Biochim Biophys Acta. 1998;1408(2-3):90–108.

31. Griese M. Pulmonary surfactant in health and human lung diseases: state of the art. Eur Respir J. 1999;13(6):1455–76.

32. Baker AD, Malur A, Barna BP, Ghosh S, Kavuru MS, Malur AG, et al. Targeted PPAR{gamma} deficiency in alveolar macrophages disrupts surfactant catabolism. J Lipid Res. 2010;51(6):1325–31.

