## Supplemental Data for "Alveolar macrophage lipid burden correlates with clinical improvement in patients with Pulmonary Alveolar Proteinosis"

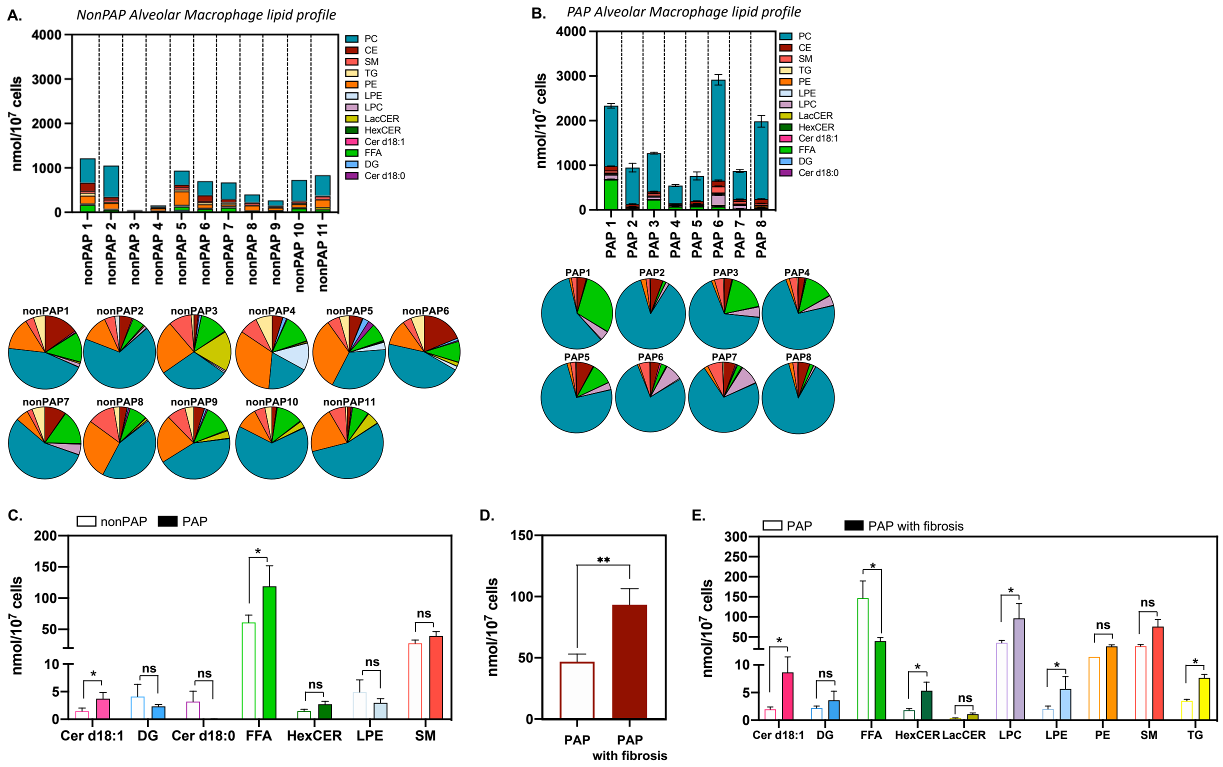


**Supplemental Figure 1. High-resolution unbiased lipidomic profiling reveals broad changes in lipid metabolism in PAP alveolar macrophages.** (A-B) Quantitative compositional profiling of lipid classes in alveolar macrophages of individual nonPAP patients (A) and PAP patients (B). (C) Total Cer d18:1, DG, Cer d18:0, FFA, HexCER, LPE, and SM content in alveolar macrophages of nonPAP and PAP patients. (D) Total CE and (E) Cer d18:1, DG, FFA, HexCER, LacCer, LPC, LPE, SM, and TG content in alveolar macrophages of PAP patients and PAP patients with fibrosis. All samples are run in duplicate or triplicate. Data are mean ± SEM. CE = cholesterol esters; Cer d18:1= ceramides; Cer d18:0= dihydroceramides; DG= diacylglycerols; FFA= free fatty acids; HexCER= hexosylceramides; LacCER= lactosylceramides; LPC= lysophosphatidylcholines; LPE= lysophosphatidylethanolamines; PC= phosphatidylcholines; PE= phosphatidylethanolamines; SM= sphingomyelins; TG= triacylglycerols. Significance was determined at $p<0.05$ by Student's $t$-test. * $p<0.05$, ** $p<0.01$, *** $p<0.001$.

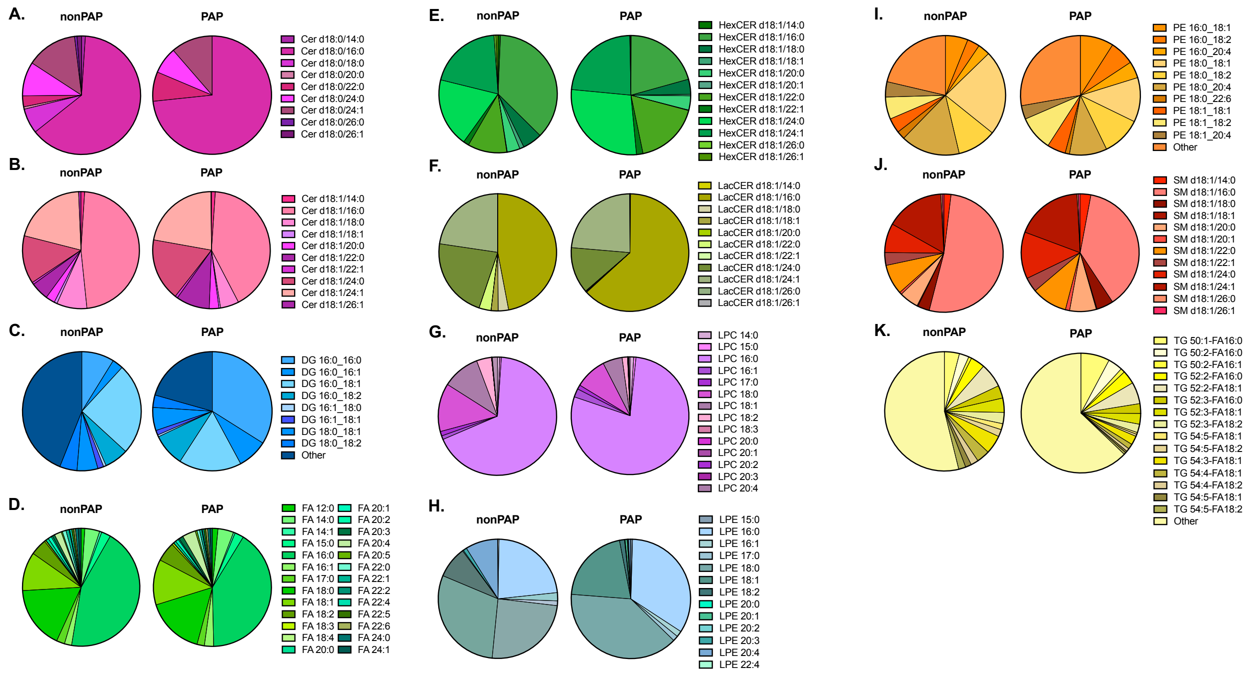


**Supplemental Figure 2. PAP alveolar macrophages display a significant shift in lipid compositional content.** Compositional analysis of Cer d18:0 (A), Cer d18:1 (B), DG (C), FA (D), HexCer (E), LacCER (F), LPC (G), LPE (H), PE (I), SM (J), and TG (K) in alveolar macrophages from nonPAP and PAP patients. All samples are run in duplicate or triplicate. Data are mean ± SEM. Cer d18:1= ceramides; Cer d18:0= dihydroceramides; DG= diacylglycerols; FFA= free fatty acids; HexCER= hexosylceramides; LacCER= lactosylceramides; LPC= lysophosphatidylcholines; LPE= lysophosphatidylethanolamines; PE= phosphatidylethanolamines; SM= sphingomyelins; TG= triacylglycerols.
